# Neural dynamics of variable grasp movement preparation in the macaque fronto-parietal network

**DOI:** 10.1101/179143

**Authors:** Jonathan A Michaels, Benjamin Dann, Rijk W Intveld, Hansjörg Scherberger

## Abstract

Our voluntary grasping actions lie on a continuum between immediate action and waiting for the right moment, depending on the context. Therefore, studying grasping requires investigating how preparation time affects this process. Two macaque monkeys (Macaca mulatta) performed a grasping task with a short instruction followed by an immediate or delayed go cue (0-1300 ms) while we recorded in parallel from neurons in the hand area (F5) of the ventral premotor cortex and the anterior intraparietal area (AIP). Initial population dynamics followed a fixed trajectory in the neural state space unique to each grip type, reflecting unavoidable preparation, then diverged depending on the delay. Although similar types of single unit responses were present in both areas, population activity in AIP stabilized within a unique memory state while F5 activity continued to evolve, tracking subjective anticipation of the go cue. Intriguingly, activity during movement initiation clustered into two trajectory clusters, corresponding to movements that were either ‘as fast as possible’ or withheld movements, demonstrating a widespread state shift in the fronto-parietal grasping network when movements must be withheld. Our results reveal how dissociation between static and dynamic components of movement preparation as well as differentiation between cortical areas is possible through population level analysis.

**Significance Statement:** Many of our movements must occur with no warning, while others we can prepare in advance. Yet, it’s unclear how planning for movements along the spectrum between these two situations differs in the brain. Two macaque monkeys made reach to grasp movements after varying amounts of preparation time while we recorded from premotor and parietal cortex. We found that the initial response to a grasp instruction was specific to the required movement, but not the preparation time, reflecting required processing. However, when more preparation time was given, neural activity achieved unique states that likely related to withholding movements and anticipation of movement, which was more prevalent in premotor cortex, suggesting differing roles of premotor and parietal cortex in grasp planning.

## Introduction

Some actions, such as reacting to a spilling cup of coffee, demand an immediate response. Others, such as waiting before a traffic light, require withholding our actions for the right moment. Most of our actions lie on the continuum between the two, and although many actions are carefully planned before they are executed (Kutas and Donchin, 1974; Ghez et al., 1997), we are often required to act with little or no warning. Various studies have examined how movements are planned and held in memory in the primate brain (Wise, 1985; Riehle and Requin, 1989), but only a few have contrasted well planned movements with situations where little to no preparation is possible (Wise and Kurata, 1989; Crammond and Kalaska, 2000; Churchland et al., 2006; Yu et al., 2009; Ames et al., 2014). None, to our knowledge, have systematically probed the transition between immediate and planned grasping movements in the behaving primate.

Delayed movement paradigms, in which actions must be withheld before they are executed, have shown that activity in premotor and parietal cortex can be used to decode and disentangle object properties and hand shapes during preparation (Baumann et al., 2009; Fluet et al., 2010; Townsend et al., 2011; Schaffelhofer et al., 2015; Schaffelhofer and Scherberger, 2016) and during movement (Menz et al., 2015). Furthermore, preparatory activity in the premotor (Churchland et al., 2006; Afshar et al., 2011) and parietal cortex (Snyder et al., 2006; Michaels et al., 2015) is correlated with reach and grasp reaction time, and perturbing this preparation state in premotor cortex delays subsequent movement (Day et al., 1989; Churchland and Shenoy, 2007; Gerits et al., 2012), a clear indication of a functional contribution to action planning.

Recent studies, made possible by the increasing implementation of large-scale sequential and parallel recordings, have employed a state space framework of population activity (for a review see Cunningham and Yu, 2014). Under this framework, the firing of each neuron represents a dimension in a high-dimensional space of all neurons where the firing of all neurons at a particular time represents a single point in the space of all potential network states. For example, preparatory activity in motor cortex acts as an initial state for subsequent movement dynamics (Churchland et al., 2012), and when reaches are cued immediately the neural population in dorsal premotor cortex (PMd) can bypass the state space achieved during delayed movements (Ames et al., 2014), suggesting that successful preparation of the same reach may be achieved through different neural trajectories. After adequate preparation time activity stabilized in the state space, while other studies suggest that premotor cortex may track time or expectation (Carnevale et al., 2015). Only analyzing the full continuum of preparation from immediate to fully planned movements can provide an understanding of the complex interaction between planning and movement. Furthermore, it has been proposed that delayed and immediate movements are controlled quite differently (Braver, 2012), a feature that has not been investigated in premotor cortex.

To address these questions, we recorded neural populations from the grasping circuit consisting of the hand area (F5) of the ventral premotor cortex (PMv) and the anterior intraparietal area (AIP) while two macaque monkeys performed a delayed grasping task, with a memory component, in which preparation time was systematically varied using 12 discrete delays (0-1300 ms). We found that the neural states achieved during longer delays were bypassed during immediately cued grasps. However, the initial trajectory was specific to each grip type, but the same regardless of delay, providing evidence that this activity may be required for successful movement. Activity in AIP stabilized during long delays, but activity in F5 was highly dynamic and well matched the subjective probability of a cue throughout the memory period. Interestingly, activity in both areas formed distinct long and short delay trajectory clusters following the go cue, demonstrating that a network-wide shift occurs when movements are withheld and executed from memory.

## Materials and Methods

### Basic procedures

Neural activity was recorded simultaneously from area F5 and area AIP in one male and one female rhesus macaque monkey (Macaca mulatta, monkeys B and S; body weight 11.2 and 9.7 kg, respectively). Animal care and experimental procedures were conducted in accordance with German and European law and were in agreement with the *Guidelines for the Care and Use of Mammals in Neuroscience and Behavioral Research* (National Research Council, 2003).

Basic experimental methods have been described previously (Michaels et al., 2015; Dann et al., 2016). We trained monkeys to perform a delayed grasping task. They were seated in a primate chair and trained to grasp a handle with the left (monkey B) or the right hand (monkey S) (Figure 1a). A handle was placed in front of the monkey at chest level at a distance of ∼26 cm and could be grasped either with a power grip (opposition of fingers and palm) or precision grip (opposition of index finger and thumb; Figure 1b insets). Two clearly visible recessions on either side of the handle contained touch sensors that detected thumb and forefinger contact during precision grips, whereas power grips were detected using an infrared light barrier inside the handle aperture. The monkey was instructed which grip type to make by means of two colored LED-like light dots projected from a TFT screen (CTF846-A; Screen size: 8” digital; Resolution 800x600; Refresh rate: 75Hz) onto the center of the handle via a half mirror positioned between the monkey’s eyes and the target. A mask preventing a direct view of the image was placed in front of the TFT screen and two spotlights placed on either side could illuminate the handle. Apart from these light sources, the experimental room was completely dark. In addition, one or two capacitive touch sensors (Model EC3016NPAPL; Carlo Gavazzi) were placed at the level of the monkey’s mid-torso and functioned as handrest buttons, preventing any premature movement of the hands. The non-acting arm of monkey B was placed in a long tube, preventing it from interacting with the handle. Monkey S was trained to keep her non-acting hand on an additional handrest button.

**Figure 1.**
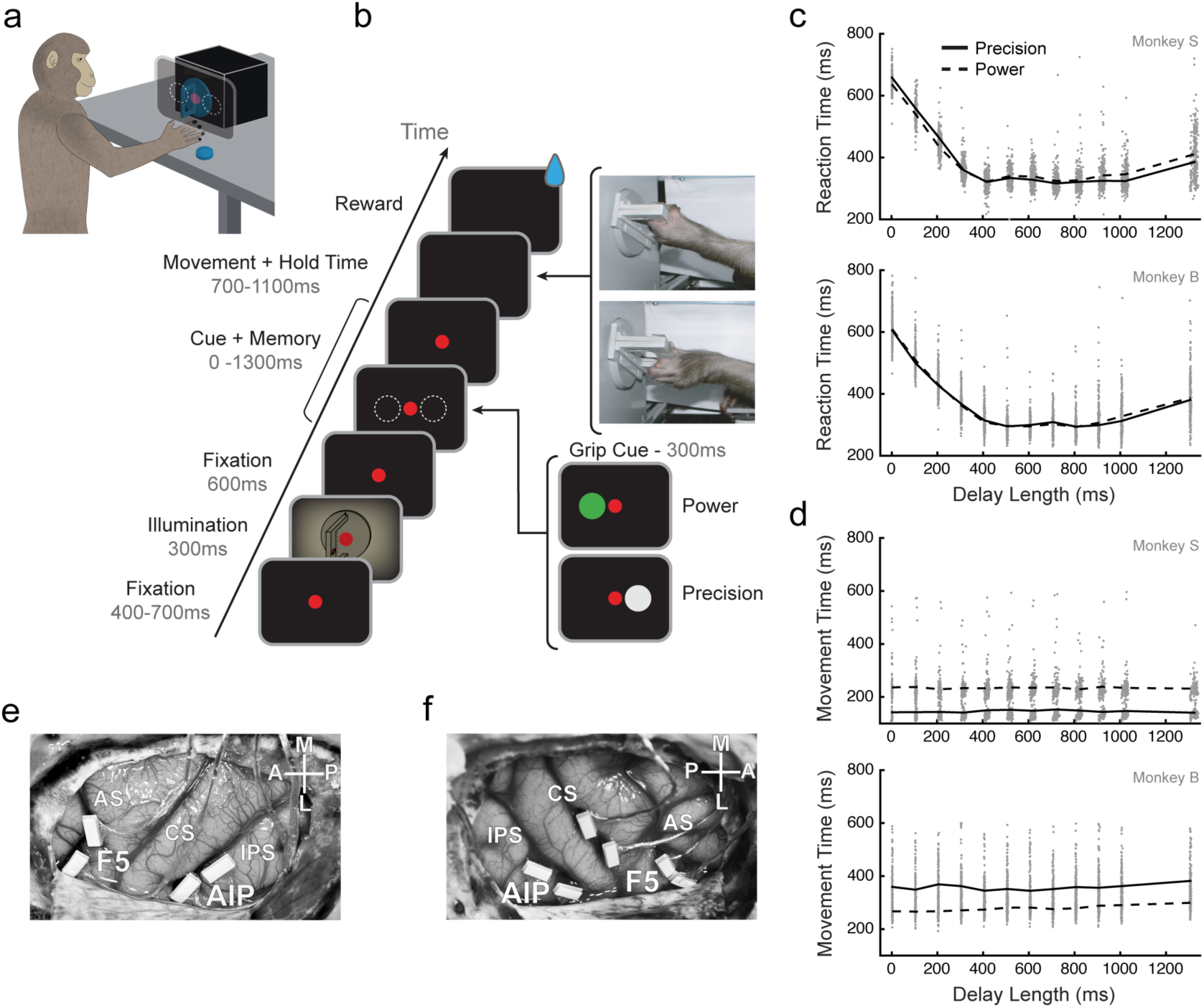
Task design, implantation, and behavior. (a) Illustration of a monkey in the experimental setup. The cues were presented on a masked monitor and reflected by a mirror such that cues appeared super-imposed on the grasping handle. (b) Delayed grasping task with two grip types (top: power grip, bottom: precision grip). Trials were presented in pseudorandom order in darkness and with the handle in the upright position. (c and d) Scatter plots of reaction time and movement time against delay length for both monkeys. The solid line represents the mean for each delay bin. (e and f) Array locations for monkey S (e) and B (f). Two arrays were placed in F5 on the bank of the arcuate sulcus (AS) and two were placed in AIP toward the lateral end of the intraparietal sulcus (IPS). In monkey B two more arrays were placed on the bank of the Central sulcus (CS), but not used in this study. The cross shows medial (M), lateral (L), anterior (A), and posterior (P) directions. Note that monkey S was implanted in the left hemisphere and monkey B the right hemisphere.

Eye movements were measured using an infrared optical eye tracker (model AA-ETL-200; ISCAN) via a heat mirror directly in front of the monkey’s head. To adjust the gain and offset, red calibration dots were shown at different locations at the beginning of each session for 25 trials that the monkey fixated for at least 2 seconds. Eye tracking and the behavioral task were controlled by custom-written software implemented in LabView Realtime (National Instruments) with a time resolution of 1 ms. An infrared camera was used to monitor behavior continuously throughout the entire experiment, additionally ensuring that monkeys did not prematurely move their hands or arms.

### Task Design

The trial course of the delayed grasping task is shown in Figure 1b. Trials started after the monkey placed the acting hand on the resting position and fixated a red dot (fixation period). The monkey was required to keep the acting hand, or both hands (monkey S), completely still on the resting position until 150 ms after the go cue. After a variable period of 400 to 700 ms two flashlights illuminated the handle for 300 ms, followed by 600 ms of additional fixation. In the cue period a second light dot was then shown next to the red one to instruct the monkey about the grip type for this trial (grip cue). Either a green or white dot appeared for 300 ms, indicating a power or a precision grip, respectively. After that, the monkey had to either react immediately or memorize the instruction for a variable memory period (also referred to as delay length). This memory period lasted for 0 to 1300 ms, in discrete memory period bins of 0, 100, 200, 300, 400, 500, 600, 700, 800, 900, 1000, or 1300 ms (i.e. the go cue could appear simultaneously with the grip cue, which was always presented for 300 ms regardless of the length of the delay). Switching off the fixation light then cued the monkey to reach and grasp the target (movement period) in order to receive a liquid reward. Monkeys were required to hold the appropriate grip for 300 ms. A failed trial occurred if the monkeys stopped fixating the central point before movement onset, moved their hand from the hand rest sensor, performed the incorrect grip, or took longer than 1100 ms to complete the movement following the go cue. Additionally, no-movement trials were randomly interleaved (8% of trials), in which a go cue was never shown and the monkey only received a reward if it maintained fixation and the hands on the hand rests for 2000 ms following the grip cue. All trials were randomly interleaved and, apart from the 300 ms handle illumination period, in total darkness.

### Surgical procedures and imaging

Upon completion of behavioral training, each monkey received an MRI scan to locate anatomical landmarks, for subsequent chronic implantation of microelectrode arrays. Each monkey was sedated (e.g., 10 mg/kg ketamine and 0.5 mg/kg xylazine, i.m.) and placed in the scanner (GE Healthcare 1.5T or Siemens Trio 3T) in a prone position. T1-weighted volumetric images of the brain and skull were obtained as described previously **(Baumann et al., 2009)**. We measured the stereotaxic location and depth orientation of the arcuate and intra-parietal sulci to guide placement of the electrode arrays.

An initial surgery was performed to implant a head post (titanium cylinder; diameter, 18 mm). After recovery from this procedure and subsequent training of the task in the head-fixed condition, each monkey was implanted with floating microelectrode arrays (FMAs; MicroProbe for Life Science) in a separate procedure. Monkey B was implanted with six electrode arrays in the right hemisphere, each with 32 electrodes (Figure 1e). Two such arrays were implanted in area F5, two in area AIP, and two in area M1. Monkey S was implanted with four FMAs in the left hemisphere and received two arrays in each area (Figure 1f). The arcuate sulcus of monkey S did not present a spur, but in the MRI a small indentation was visible in the posterior bank of the arcuate sulcus, about 2 mm medial to the knee, which we treated as the spur. We placed both anterior FMAs lateral to that mark. FMAs consisted of non-moveable monopolar platinum-iridium electrodes with initial impedances ranging between 300 and 600 kΩ at 1 kHz measured before implantation and verified in vivo. Lengths of electrodes were between 1.5 and 7.1 mm.

All surgical procedures were performed under sterile conditions and general anesthesia (e.g., induction with 10 mg/kg ketamine, i.m., and 0.05 mg/kg atropine, s.c., followed by intubation, 1–2% isofluorane, and analgesia with 0.01 mg/kg buprenorphene). Heart and respiration rate, electrocardiogram, oxygen saturation, and body temperature were monitored continuously and systemic antibiotics and analgesics were administered for several days after each surgery. To prevent brain swelling while the dura was open, the monkey was mildly hyperventilated (end-tidal CO_2_, ∼30 mmHg) and mannitol was kept at hand. Monkeys were allowed to recover fully (∼2 weeks) before behavioral training or recording experiments commenced.

### Neural recordings and spike sorting

Signals from the implanted arrays were amplified and digitally stored using a 128 channel recording system (Cerebus, Blackrock Microsystems; sampling rate 30 kS/s; 0.3-7500Hz hardware filter; see Supplementary Methods). Data were first filtered using a median filter (window-length: 3ms) and the result subtracted from the raw signal, corresponding to a nonlinear high-pass filter. Afterwards, the signal was low-pass filtered with a non-causal Butterworth filter (5000 Hz; 4^th^ order). To eliminate movement noise (i.e., common component induced by reference and ground), PCA artifact cancellation was applied for all electrodes of each array (Musial et al., 2002; Dann et al., 2016). In order to ensure that no individual channels were eliminated, PCA dimensions with any coefficient greater than 0.36 (with respect to normalized data) were retained. Spike waveforms were extracted and semi-automatically sorted using a modified version of the offline spike sorter Wave_clus (Quiroga et al., 2004; Kraskov et al., 2009).

Units were classified as single- or non-single unit, based on five criteria: (1) the absence of short (1–2 ms) intervals in the inter-spike interval histogram for single units, (2) the homogeneity and SD of the detected spike waveforms, (3) the separation of waveform clusters in the projection of the first 17 features (a combination for optimal discriminability of principal components, single values of the wavelet decomposition, and samples of spike waveforms) detected by Wave_clus, (4) the presence of well known waveform shapes characteristics for single units, and (5) the shape of the inter-spike interval distribution.

After the semiautomatic sorting process, redetection of the average waveforms (templates) was done in order to detect overlaid waveforms (Gozani and Miller, 1994). Filtered signals were convolved with the templates starting with the biggest waveform. Independently for each template, redetection and resorting was run automatically using a linear classifier function (Matlab function: classify). After the identification of the target template, the shift-corrected template (achieved by up and down sampling) was subtracted from the filtered signal of the corresponding channel to reduce artifacts for detection of the next template. This procedure allowed a detection of templates up to an overlap of 0.2 ms. Unit isolation was evaluated again as described before to determine the final classification of all units into single- or multi-units. Units were only classified as single if they unambiguously met the five criteria.

### Data preprocessing

Although units were classified as single- or multi-units, all recorded units were used for all analyses. A detailed list of data set information can be found in Table 1. After spike sorting, spike events were binned in non-overlapping 1 ms windows. For individual unit plotting (Figure 2), spike trains were smoothed with a Gaussian window (σ = 50 ms), but for all analyses spike trains were further reduced to a set of latent dimensions (see next section). Data were aligned to two events, the presentation of the grip cue and movement onset, i.e. the time when the monkey’s hand left the handrest button. The cue alignment proceeded from 200 ms before cue onset until the go cue, and the movement onset alignment from movement onset minus the median reaction time for each delay condition until 400 ms after movement onset. These two alignments were combined to produce a continuous signal. In this case the two signals were simply concatenated in time. Average firing rates were then calculated by averaging over all trials of the same condition.

**Table 1.** Trial counts, performance, and number of units recorded for all data sets.

|  | Trial<br>Count | Correct<br>Performance | Units Recorded<br>in F5 | Units Recorded<br>in AIP |
| --- | --- | --- | --- | --- |
| B1 | 485 | 91% | 65 | 29 |
| B2 | 685 | 96% | 88 | 35 |
| B3 | 586 | 96% | 43 | 25 |
| B4 | 814 | 96% | 64 | 28 |
| B5 | 775 | 96% | 46 | 19 |
| B6 | 745 | 97% | 72 | 33 |
| Mean: | 682 | 95.3% | 63.0 | 28.2 |
| S1 | 502 | 98% | 124 | 134 |
| S2 | 514 | 97% | 136 | 148 |
| S3 | 571 | 97% | 142 | 137 |
| S4 | 658 | 99% | 121 | 97 |
| S5 | 590 | 99% | 115 | 104 |
| S6 | 546 | 98% | 156 | 165 |
| Mean: | 564 | 98.0% | 132.3 | 130.8 |

**Figure 2.**
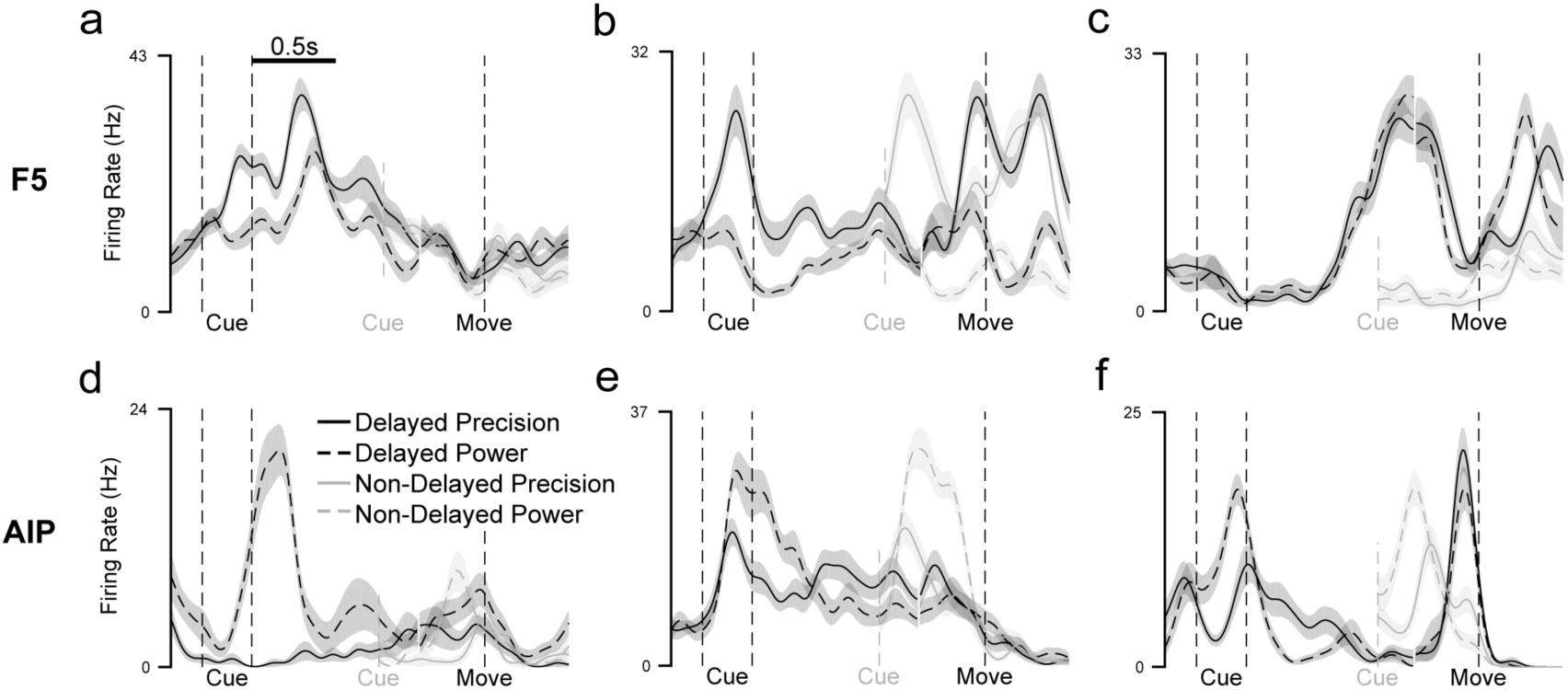
Example average firing rate curves of single-units for delayed (1300 ms) vs. non-delayed (0 ms) grasps. (a-c) Example single-units from area F5 of monkey B showing (a) a completely suppressed cue response during non-delayed grasps, (b) an identical cue response for either delay, (c) differing movement period activity between delayed and non-delayed grasps. (d-f) Similar single-unit examples from AIP of monkeys B and S. Delayed data were aligned to two events, grip cue onset and movement onset and are separated by a gap, which marks the go cue. Non-delayed data were only aligned to movement onset. Dotted gray line represents approximate time of cue onset and go cue for non-delayed grasps. The cue was always presented for 300 ms regardless of delay. Curves and shaded bands represent mean and standard error of the mean, respectively.

### Dimensionality reduction

In order to extract a set of smooth single-trial neural trajectories in our neural populations we applied Gaussian Process Factor Analysis (GPFA; Yu et al., 2009) to all neurons of both areas over all successful trials from 200 ms before cue onset to 400 ms after movement onset for each recording session separately. Performing a single dimensionality reduction over both areas allows a direct comparison of each area’s contribution to the common signals. Units within each session were recorded simultaneously across both areas. GPFA is similar to factor analysis in that it finds an explanatory set of orthogonal dimensions based on the covariance structure between units that is a linear combination of binned neural data. However, in GPFA, each dimension denoises data with a Gaussian smoothing kernel of unique width learned from the data. For our GPFA analysis, neural spiking data on single trials were binned into 50 ms bins and square-rooted before being transformed through linear combination into 10 latent dimensions. Units with an average firing rate less than 1 Hz were discarded before the analysis. These 10 dimensions, each based on an individual smoothing kernel, were further orthonormalized to produce a set of 10 orthogonal dimensions, each containing a combination of all smoothing kernels. Cross-validation procedures were undertaken to determine the optimal number of latent dimensions (Yu et al., 2009). Beyond 10 latent dimensions very little shared variance was explained by further addition of dimensions (<3% per dimension), and visualization of these dimensions showed almost no modulation.

Since GPFA was carried out across both recorded areas simultaneously, to identify the specific contribution of each area to each latent dimension the neural data of each area were separately transformed into the previously found latent dimensions. In more detail, the smoothing kernels and transformation matrix found through GPFA over both areas was used to independently transform the binned neural data from each area into the 10 latent dimensions, giving a representation of each latent dimension as a linear combination of data from either F5 or AIP. For most analyses the extracted single trials were then cut into two alignments (previous section) and averaged over all trials of the same condition. In general. at the boundary of alignments the signals matched very well to each other, showing almost no jumps in activity.

### Distance analysis

In order to find the neural distance between two conditions over time, we calculated the minimum Euclidean distance (point-to-curve distance) between the two trajectories in the space of the 10 latent dimensions extracted through GPFA separately for each area. Three versions of this analysis were performed. For the distance in Figure 4a, we iterated through all time points on delayed trajectory (in steps of 50 ms) and calculated the Euclidean point-to-curve distance from the delayed (1000 ms) trajectory to the non-delayed (0 ms) trajectory, where the point-to-curve distance is the minimum distance from a specific time point on the delayed trajectory to all points on the non-delayed trajectory. Minimum distance, as a conservative measure, was used in order to overcome the different time courses of the conditions being compared. Small distances indicate that the two trajectories achieve a similar point in neural space at some point in time, while large distances indicate that the two trajectories do not pass through a similar point in the high dimensional space. Euclidian distances were normalized by the square root of the number of neurons in order to make spaces with different number of neurons comparable.

**Figure 3.**
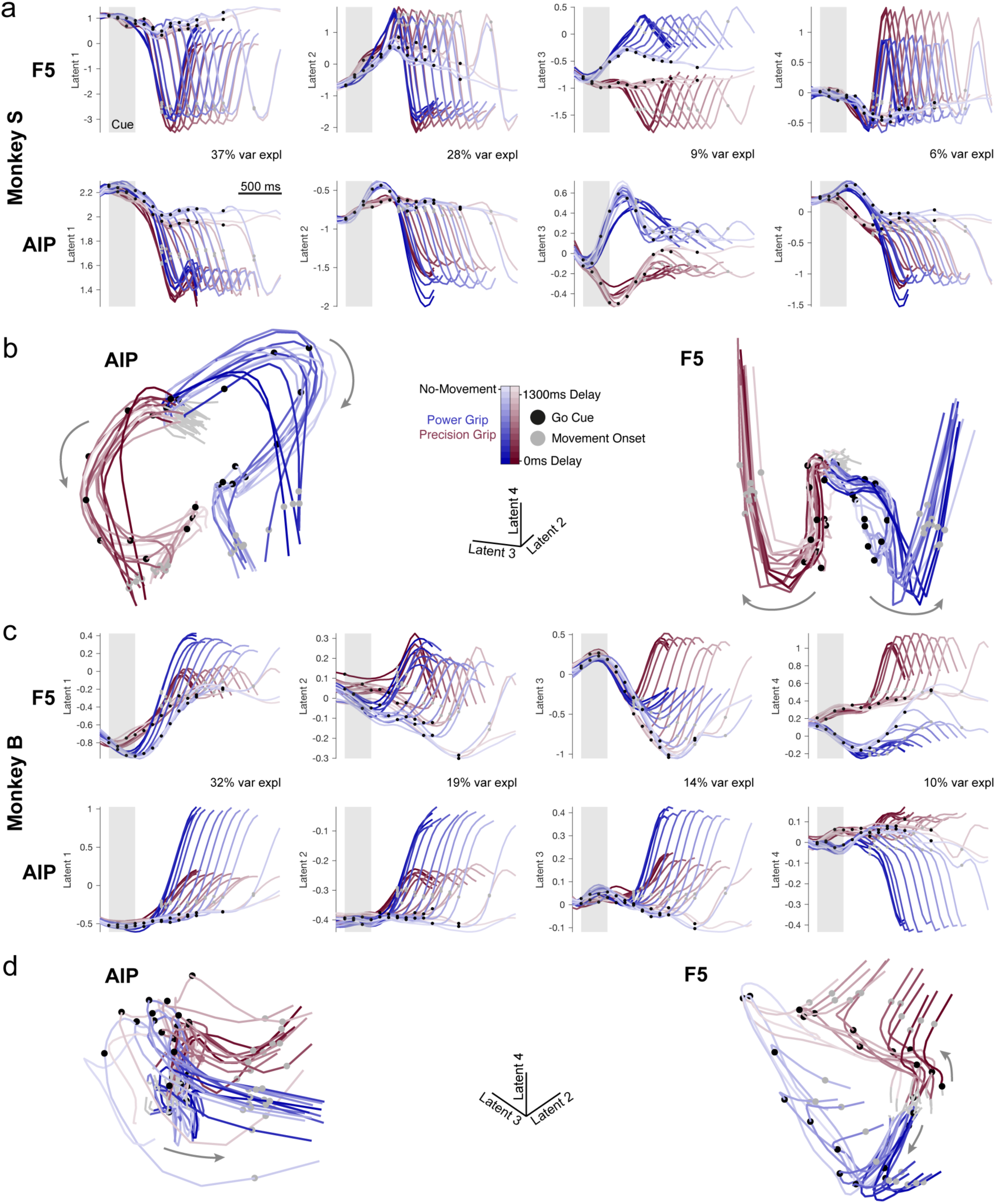
Low-dimensional latent space trajectories of F5 and AIP. Population data of all conditions were projected into a 10 dimensional latent space as determined by GPFA. (a) A single session trial-averaged example from monkey S is shown for the first 4 latent dimensions (S4). Trajectories begin 100 ms before the grip cue and end 400 ms after movement onset. (b) A 3D plot of the second to fourth latent dimensions plotted from 100 ms before cue onset to 50 ms after movement onset. (c-d) same as (a-b) for a single session from monkey B (B2). Gray arrows show the flow of time.

**Figure 4.**
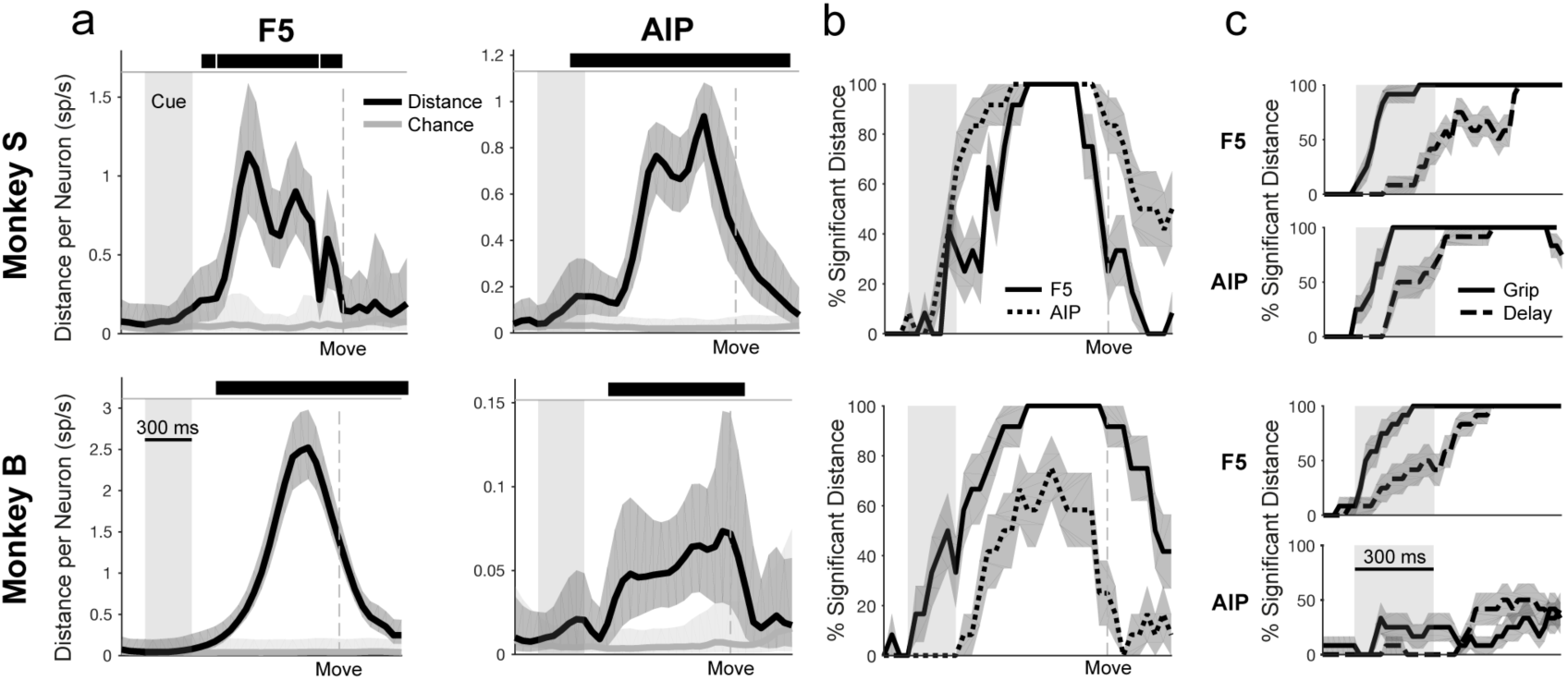
Point-to-curve distance between delayed (1000 ms) and non-delayed (0 ms) trajectories. (a) Minimum Euclidian distance in the latent space between each time point on the delayed trajectory (in steps of 50 ms) and the entire non-delayed trajectory over time for 2 example data sets (B2-Power, S3-Power) from both areas and monkeys. The black line represents the minimum point-to-curve distance between the delayed and non-delayed trajectory, while the gray lines represent the chance level (Materials and Methods). Black bars along the top of plots denote times when the distance is significantly greater than chance level (Bootstrapping procedure with 1000 resamples, p = 0.05, Cluster-based permutation test; Materials and Methods). Error bars represent the 5^th^ and 95^th^ percentiles of the distances generated by the bootstrapping procedure. (b) Fraction of significant distances over all data sets and grip types (6 data sets x 2 grip types). Error bars represent the standard error of the mean over data sets and grip types. (c) Difference in onset of grip and delay separation over all data sets and grip types (6 data sets x 2 grip types) at a higher temporal resolution (20 ms bins).

For the distance analysis in Figure 4c, GPFA was recalculated on a smaller portion of the data (200 ms before cue onset to 800 ms after) with a shorted bin width of 20 ms. Distance was then calculated as before between the delayed and non-delayed trajectories. In addition, to determine when grip information becomes present in the population, distance between the delayed trajectories (1000 ms) of each grip type was calculated in the same manner.

For the distance analysis in Figure 5, the Euclidean distance was calculated between all pairs of time points on the same trajectory (no-movement) and used in conjunction with the bootstrapping procedure (next section) to determine if two points significantly differed.

**Figure 5.**
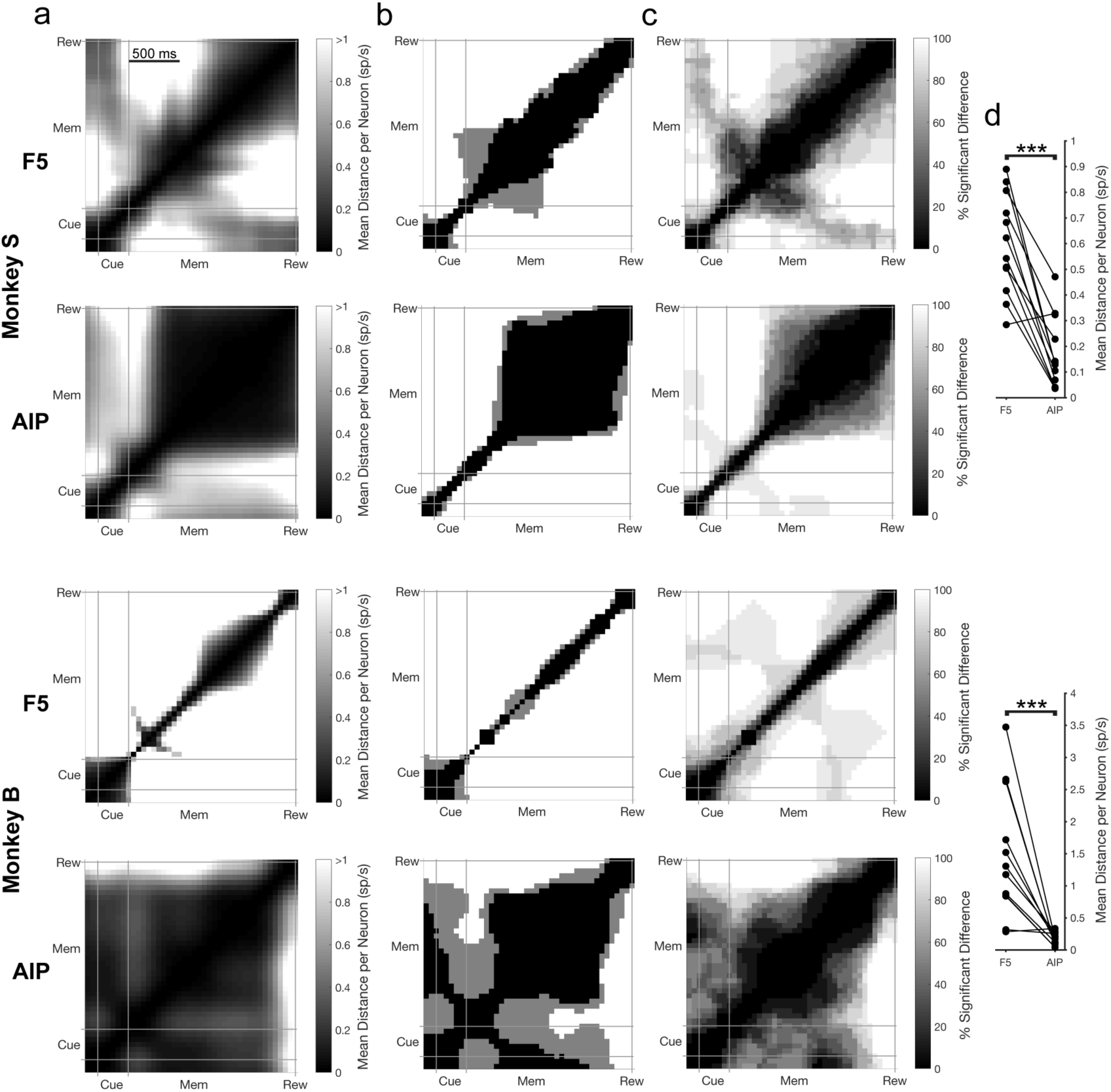
Neural trajectory stability over the course of no-movement trials. (a) Mean Euclidean distance in the latent space for the no-movement trials between all pairs of time points over both grip types for example data sets in each monkey (sessions B5, S6). For each pair of time points, distance results were tested for a significant difference using a bootstrapping procedure (10000 resamples in steps of 50 ms, p = 0.01). The abbreviations Cue, Mem, and Rew, correspond to the cue, memory, and reward epochs, respectively. All plots are clipped at 1 sp/s for visualization. The times where a significant difference was found (in no conditions, one grip type, or both grip types) are shown in (b). (c) Percentage of time points showing a significant difference over all data sets and grip types (6 data sets x 2 grip types) of each monkey separately. (d) Mean distance between all time points during the stable portion of the memory period (600 ms – 1800 ms after cue onset) for all individual data sets and grip types (6 data sets x 2 grip types) across areas and paired according to recording session. Stars indicate a significant difference (Wilcoxon sign-rank test, p < 0.001).

### Bootstrap procedure

In order to gain an estimate of underlying trial-to-trial variability, we performed a bootstrap analysis. This procedure was in general the same, but with slight variations for the different distance analyses presented above. We resampled trials from each condition randomly, with replacement, of the same size as the number of recorded trials in that condition. We then constructed average firing rates for each condition and carried out the appropriate distance analysis as described above (e.g., minimum distance between delayed and non-delayed trajectory). This resampling was done 1000 times, producing a distribution of distances.

To obtain an estimate of how much distance is expected between trajectories by chance, we carried out another resampling in which a trajectory was resampled from itself to determine its underlying variability. Trajectories were resampled once with the number of trials observed in that condition, and once using the number of trials recorded in the other trajectory in the comparison, then the Euclidean distance was calculated as described in the previous section.

To determine when the observed distance distribution was significantly greater than the self-sampled distribution, we used a cluster-based permutation test (CBPT; Maris and Oostenveld, 2007). Briefly, we used a modification of the original test that evaluates the area under the receiver operator characteristic curve (AUC) between the distance distribution and the self-sampled distribution over all time points and extracts clusters (consecutive time segments) of activity whose AUC exceeds a predefined threshold (α = 0.1), then the absolute AUCs within each cluster were summed to produce cluster-level statistics. To generate a chance-level distribution from which the cluster-level statistics could be calculated, trials were randomly partitioned between the two conditions and the AUC and clustering redone (1000 partitions). From every partition the largest cluster was used to generate a largest chance cluster distribution. Cluster-level statistics were calculated by comparing the real cluster-levels against the largest chance cluster distribution. Real clusters were considered significant if they exceeded 95% of all largest chance cluster values corresponding to a p = 0.05. In this way, sensitivity to short or small time-scale differences is greatly reduced, but the overall false-alarm rate across time points remains below the designated p-value. This analysis allowed us to determine when an observed distance was significantly greater than the distance expected if two trajectories were generated from the same underlying distribution.

For chance analyses in Figure 5, resampling of trials was carried out 10000 times, with replacement, for each condition and data set. For each of the 10000 resampling steps the same trajectory was resampled twice, termed ***p*** and ***p***’. Then, for every pair of time points (*t*_1_ and *t*_2_), the resampled distance along the first trajectory *d* = *d*(***p***(*t*_1_), ***p***(*t*_2_)) was compared to the two inter-trajectory distances at time *t*_1_ and *t*_2_: *d*_1_ = *d*(***p***(*t*_1_), ***p***’(*t*_1_)) and *d*_2_ = *d*(***p***(*t*_2_), ***p***’(*t*_2_)). We determined the percentile of resamples (across all 10000) for which the along-trajectory distance *d* exceeded both inter-trajectory distances: *d* > *max*(*d*_1_, *d*_2_). This percentile determined a specific p-value for each time pair *t*_1_, *t*_2_. The resampled distance, *d*, was then considered significant if p < 0.01. In this way, the significance level was dependent on which time points were compared along the trajectory, establishing a conservative estimate of the underlying trial-to-trial variability.

### Hazard rate

To classify the temporal evolution of activity during the memory period, the mean firing rate of each latent dimension for the no-movement condition from cue onset until reward onset was fit with an anticipation function, which can be described as the conditional probability that a movement will be required at a given moment, given that it has not occurred until this point. This type of anticipation has been termed the hazard rate, and we present it here precisely as in Janssen and Shadlen **(2005)**. The hazard rate can be expressed as

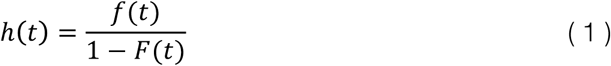

where *f*(*t*) is the probability that a go cue will come at a given time after cue onset, and *F*(*t*) is the cumulative distribution, 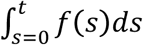

As in Janssen and Shadlen (2005), to obtain an estimate of the monkey’s internal representation of anticipation we calculate ‘subjective anticipation’ based on the assumption that the animal is uncertain about time and that this uncertainty scales with time since an event. Therefore, before calculating hazard rate we smoothed our probability density function, *f*(*t*), with a normal distribution where standard deviation is proportional to elapsed time.

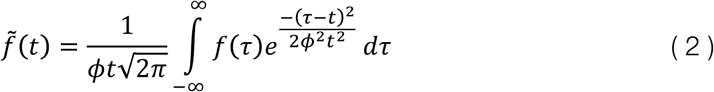

The coefficient of variation, *Φ*, is a Weber fraction under the assumption that the experience of elapsed time carries uncertainty that is proportional to the true duration (Weber’s Law). For all analyses we used a value of 0.26, as has been calculated from behavioral experiments and used previously (Leon and Shadlen, 2003; Janssen and Shadlen, 2005). To obtain the final subjective anticipation function, 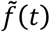 was then substituted into Eq. 1, along with its cumulative distribution, 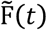.

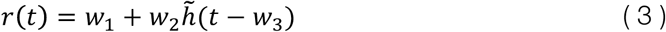

All fitting procedures were performed by fitting Eq. 3 to the average activity of each latent dimension over both areas, where *w* are constant terms obtained during the fitting procedure (Matlab function: *fit*), and 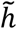 is Eq. 2 substituted into Eq. 1.

### Clustering analysis

To evaluate whether or not delay trajectories leading up to movement onset clustered in a distinct way, we calculated the Euclidean distance between all pairs of linearly spaced delays (0-1000 ms, in steps of 50 ms) in the 10 latent dimensions determined by GPFA and looked for community structure (i.e. distinct clusters of similar value) in the resulting distance matrix. We employed a well-known modularity analysis that iteratively finds non-overlapping groups of conditions that minimizes the within-group distance between conditions and maximizes the between-group distance (Newman, 2004; Reichardt and Bornholdt, 2006) with a gamma sensitivity of 0.75. Each distance matrix was normalized to the maximum value over all time and subtracted from a matrix of ones in order to prepare them for analysis. Using this analysis, the number of clusters obtained is purely data-driven and not specified by the experimenter. To ensure that the found structure was not due to chance, we randomly permuted the distance matrix (1000 permutations, while conserving matrix symmetry) and compared the modularity index Q between the empirical and permuted data. The percentile of instances where the permuted distribution values exceeded the empirical value corresponds to the p-value.

## Results

### Task and behavior

To investigate the continuum of grasp movement preparation, we trained two macaque monkeys (B and S) to perform a delayed grasping task, with a memory component, in which the amount of preparation time was systematically varied between non-delayed (0 ms) and a long delay (1300 ms) in 12 distinct increments (Materials and Methods). Monkeys fixated a central point (red), received a grip cue (300 ms) corresponding to either precision (white) or power grip (green), and were cued to perform this grip following a variable delay when the central fixation point turned off (Figure 1a-b). The performance of both monkeys was high, correctly completing trials after receiving grip information 95% and 98% of the time for monkeys B and S, respectively (Table 1). In addition to the normal task, we also randomly inserted no-movement trials to ensure that monkeys waited for the go cue before acting. Both monkeys completed these trials successfully (monkey B: 100%; monkey S: 97.7%).

In addition to number of correctly executed trials, reaction times (RTs) and movement times (MTs) of the monkeys provided useful insight into the performance of the task. RT decreased steadily with increasing amounts of preparation (Rosenbaum, 1980), approaching a minimum after approximately 400 ms of preparation (Figure 1c), well in line with previous findings (Churchland et al., 2006). RT increased slightly for the longest delay. For monkey S, MT did not correlate with length of the delay period (Figure 1d, p = 0.9), indicating that although RT was slower for short delays, movements were only initiated once they were fully prepared. In monkey B there was a small positive correlation between delay and MT (Figure 1d, r = 0.11). Movement kinematics were likely similar regardless of delay, since the variability in mean movement times between different delay lengths were extremely small. The standard deviations in mean movement times (Monkey S, precision grip: 3.5 ms SD, power grip: 1.8 ms SD; Monkey B, precision grip: 14.2 ms SD, power grip: 10.8 ms SD) provide evidence that the kinematics of the movements did not vary between delays, especially for monkey S. The number of errors showed no clear relationship to the length of the delay period, and the number of errors was extremely low, providing evidence that the monkeys could complete all conditions equally well.

### Neural responses

We recorded six sessions of each monkey using floating microelectrode arrays for a total of 128 channels (64 in each area) simultaneously in F5 and AIP (Figure 1e,f) and single- and multi-unit activity was isolated (Materials and Methods). There were significantly more units recorded in area F5 of monkey B than in AIP (Paired t-test, p < 0.001), while there was no significant difference for monkey S (Paired t-test, p = 0.81). For individual session information see Table 1. For all analyses we pooled single- and multi-units together (mean recorded per session: 75 single and 102 multi). We evaluated grip type tuning in both areas to ensure that the task successfully elicited task-related tuning. The average percentage of units tuned for grip type during the 200 ms following cue onset was 29% in F5 and 29% in AIP, 28% and 26% in the 200 ms preceding go cue, and 55% and 45% in the 200 ms following movement onset (t-test, p < 0.05), conservatively measured only for movements with a distinct memory period (i.e. ≥500 ms delay). Amounts of grip tuning were very similar between monkeys and to previous studies of both F5 and AIP (Lehmann and Scherberger, 2013; Michaels et al., 2015; Schaffelhofer et al., 2015), confirming their involvement in grasp coding.

If the brain areas we investigated were specifically coding task-related visual features, we would expect similar responses to the grip cue regardless of whether grasps were cued immediately or not. Conversely, if single units were related to execution of the correct motor plan, we should observe similar neural responses during movement regardless of when go cues were presented. Interestingly, a wide variety of mixed activity patterns were present in both areas (Figure 2). In many cases the initial cue response was suppressed when the go cue appeared concurrently with the grip information (Figure 2a,d), while in other cases the initial cue response was present regardless of delay (Figure 2b,e). Other interesting responses were observed, such as a peak in activity during the memory period (Figure 2c), and activity during the movement period which differed between delayed and non-delayed grasps (Figure 2c,f). All of these diverse types of responses were present in both F5 and AIP. The broad variety of unit responses reveals a complex interaction between differing amounts of preparation, making strict categorization of individual neurons difficult.

### Visualizing the population response

An alternative approach to categorizing single units is the state space framework, in which all units are considered as a high-dimensional space in which the firing of each unit represents one dimension. In order to visualize the complex interactions between planning and movement, we projected the population activity of all units across both areas for all trials into a lower dimensional space of 10 latent dimensions using Gaussian Process Factor Analysis (GPFA; Materials and Methods). These 10 latent dimensions well captured the variance of both areas. Once the latent dimensions were found, the activity of each area was independently projected into these dimensions in order to compare the contribution of each area. Figure 3a,c shows the neural trajectories of exemplar data of each monkey (sessions B4, S2) from 100 ms before grip cue onset to 400 ms after movement onset.

In both monkeys the first dimension was a mostly condition-independent movement signal, especially large in F5, a feature observed previously in motor cortex (Kaufman et al., 2016). The other dimensions show varying levels of grip-specific cue responses, delay- or grip-specific memory responses, and strong movement activity. Particularly interesting is latent 3 in Figure 3a and latent 4 in Figure 3c, which showed in both monkeys sustained grip selectivity through memory into movement. Plotting latents 2-4 against each other revealed other features (Figure 3b,d, 100 ms before cue onset to 50 ms after movement onset). Trajectories began in a tight cluster at grip cue onset and remained overlapped for the initial response (200-300 ms) regardless of delay, but specific to each grip type. The trajectories for longer delays continued to evolve for hundreds of milliseconds, but the short delays proceeded to movement onset, bypassing the part of the space achieved by long delays. Interestingly, while activity in AIP congregated in a stable state 500-600 ms after the grip cue, activity in F5 continued to evolve for the entire memory period, never congregating in an area of low variability. Finally, for each grip type short and long delays grouped into two clusters during movement initiation (Figure 3b, AIP; Fig 3d, F5).

### Unique memory state for delayed grasping movements

As we saw in Figure 3, unique memory states were traversed by the neural trajectory during trials with long delays. To test this possibility statistically, we used a continuous distance analysis (Materials and Methods). We measured the minimum Euclidean distance (known as point-to-curve) between each time point on the trajectory of a delayed condition (1000 ms delay condition in steps of 50 ms) and the entire non-delayed trajectory (0 ms delay condition). This was done for the 10 latent dimensions of each area to determine which points in the state space were traversed by both conditions and which were unique to longer delayed movements, separately for each recording session and each grip type. After the cue, distance between delayed and non-delayed trajectories rose and remained significantly above chance level until around movement onset or later in example data sets of both areas and monkeys (Figure 4a; sessions B2, S3; Bootstrapping procedure with 1000 resamples, p < 0.05, cluster-based permutation test; Materials and Methods). Over all grip types and data sets the same effect is present (Figure 4b), showing that distance between the trajectories was most prevalent until shortly before movement onset. The amount of divergence between the delayed and non-delayed trajectories was very similar in F5 and AIP, indicating that when grasps are cued without a delay the neural population of both areas bypass the states achieved by longer delays. Performing the same analysis on the full neural space without dimensionality reduction produced similar results (data not shown).

As mentioned earlier, it appeared in Figure 3 that the difference between grip types was present before the difference between delays. In other words, the effect of the grip cue appeared before the effect of the go cue. To test this, we repeated the distance analysis with a finer time resolution around cue onset (GPFA using steps of 20 ms) and additionally tested the Euclidean distance between grip conditions (Figure 4c, Materials and Methods). Comparing the first onset of significance between delay and grip effects for each data set separately revealed that grip separation consistently appeared before delay separation in both areas and monkeys (Wilcoxon sign-rank test, F5 monkey S, p < 0.001; AIP monkey S, p < 0.001; F5 monkey B, p = 0.003; AIP monkey B, p = 0.016). On average across monkeys and areas, grip separation occurred 128 ms after cue onset and delay separation occurred 352 ms after cue onset.

Taken together, these results provide evidence that large portions of the state space that are traversed after the first ∼300 ms do not seem to be necessary for successfully executing grasping movements, and the activity in the first ∼300 ms likely represents unavoidable processing.

### Static and dynamic memory states

Given that the trajectories of delayed and non-delayed grasps only overlap for the first ∼300 ms of preparation, what is the function and dynamics of the memory period activity? A striking feature of the visualization in Figure 3 was that the F5 activity continually evolved throughout the course of the memory period, while activity in AIP congregated in an area of low variability. To analyze when and if the neuronal trajectory of the two areas stabilized, we systematically compared the Euclidean distance between all pairs of time points along the trajectories for the no-movement trajectories (Figure 5a, example data sets S6 and B5). Dynamic activity should appear as large distances between trajectories everywhere except the diagonal (points close in time), while static activity should appear as a ‘block’ of activity with a small distance between trajectories.

The strongest differences occurred shortly after cue onset and near reward. Most remarkably, the neuronal trajectory during the memory period in F5 continuously and uniformly progressed in the absence of behavioral events. On the contrary, the neuronal trajectory in AIP stabilized 200-300 ms after cue offset. The effect becomes clearer when visualizing the time points that significantly differed (Figure 5b, Materials and Methods), showing a stereotypical ‘block’ pattern in AIP and also visible over all data sets (Figure 5c). Taking the average distance between all time points during the portion of the memory period unaffected by cue or reward (600 ms – 1800 ms after cue onset) showed a significantly more dynamic representation in F5 than AIP (Figure 5d; Wilcoxon signed-rank test, p < 0.001). Similar results were obtained using the full neural space (data not shown). These results indicate a considerably different code at the population level in AIP and F5.

It is also important to consider that the probability of having to perform a movement did not remain constant, since the probability of being in the no-movement condition increased with time spent in the memory period. Therefore, could it be that the dynamic nature of the memory period in F5 is due to the change in necessity of the motor plan. To rule out this possibility, we repeated the current analysis on data of a similar experiment in which movements were required in all conditions (Michaels et al., 2015). We found that the same inter-area difference reported here were present (Figure 6), lending support to the observed dissociation between areas.

**Figure 6.**
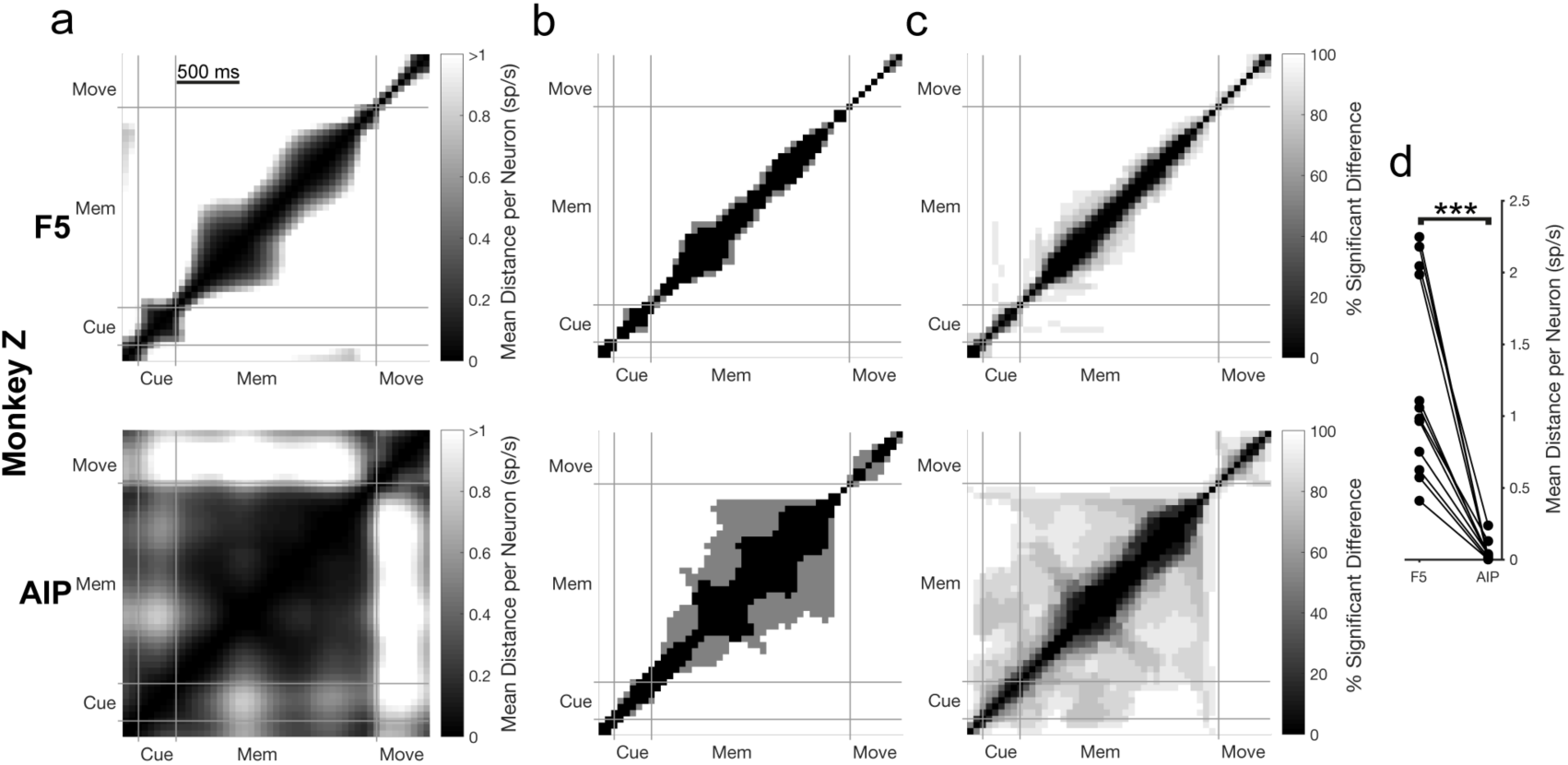
Neural trajectory stability over the course of instructed trials for an additional experiment. Same layout as Figure 5. (a) Mean Euclidean distance in the latent space for the Instructed trials between all pairs of time points over both grip types for an example data set in monkey Z. For each pair of time points, distance results were tested for a significant difference using a bootstrapping procedure (10000 resamples in steps of 50 ms, p = 0.01). The abbreviations Cue, Mem, and Move, correspond to the cue, memory, and movement epochs, respectively. All plots are clipped at 1 sp/s for visualization. The times where a significant difference was found are shown in (b). (c) Percentage of time points showing a significant difference over all data sets and grip types (6 data sets x 2 grip types). (d) Mean distance over the stable portion of the memory period (600 ms after cue onset – go cue) for all individual data sets and grip types (6 data sets x 2 grip types) across areas and paired according to recording session. Stars indicate a significant difference (Wilcoxon sign-rank test, p < 0.001). As described in Michaels et al. (2015), monkey Z performed a similar task to the current study (6 data sets x 2 grip types, Instructed condition). The same grip types were cued and the memory period was also variable. However, all trials resulted in movement, regardless of condition. Therefore, if the dynamic nature of the memory period observed in the present experiment were due only to the changing expectation of having to execute a movement over the course of the trial or the deterioration of a motor plan, we should observe stable activity. Yet, in this additional experiment the highly time dependent nature of the memory period activity un F5 is maintained, suggesting that this variability is not due to the varying chance of subsequent movement, but represents features of the examined areas.

### Memory period dynamics

Given the dynamic nature of activity during the memory period, does this activity follow any predictable pattern? As mentioned earlier, some units appeared to change their activity strictly during the memory period (Figure 2c), even in the absence of behavioral cues. The observed pattern appears similar to the hazard rate, which in the current experiment is the probability of a go cue occurring at any moment, given that the go cue has not appeared yet (Janssen and Shadlen, 2005). The form of the hazard rate during no-movement trials and corresponding subjective anticipation function, which takes the monkey’s uncertainty about time into account (Materials and Methods), is shown in Figure 7a. We fit the average activity of each latent dimension (over both areas) to subjective anticipation. The best fitting dimension per data set had an average adjusted R-square of 0.73 for monkey S and 0.88 for monkey B, indicating that anticipation may be significantly represented (mean time shift: -11 ms, w_3_ in Eq. 3). Example data sets are shown in Figure 7b,e (data from session S2 and B4).

**Figure 7.**
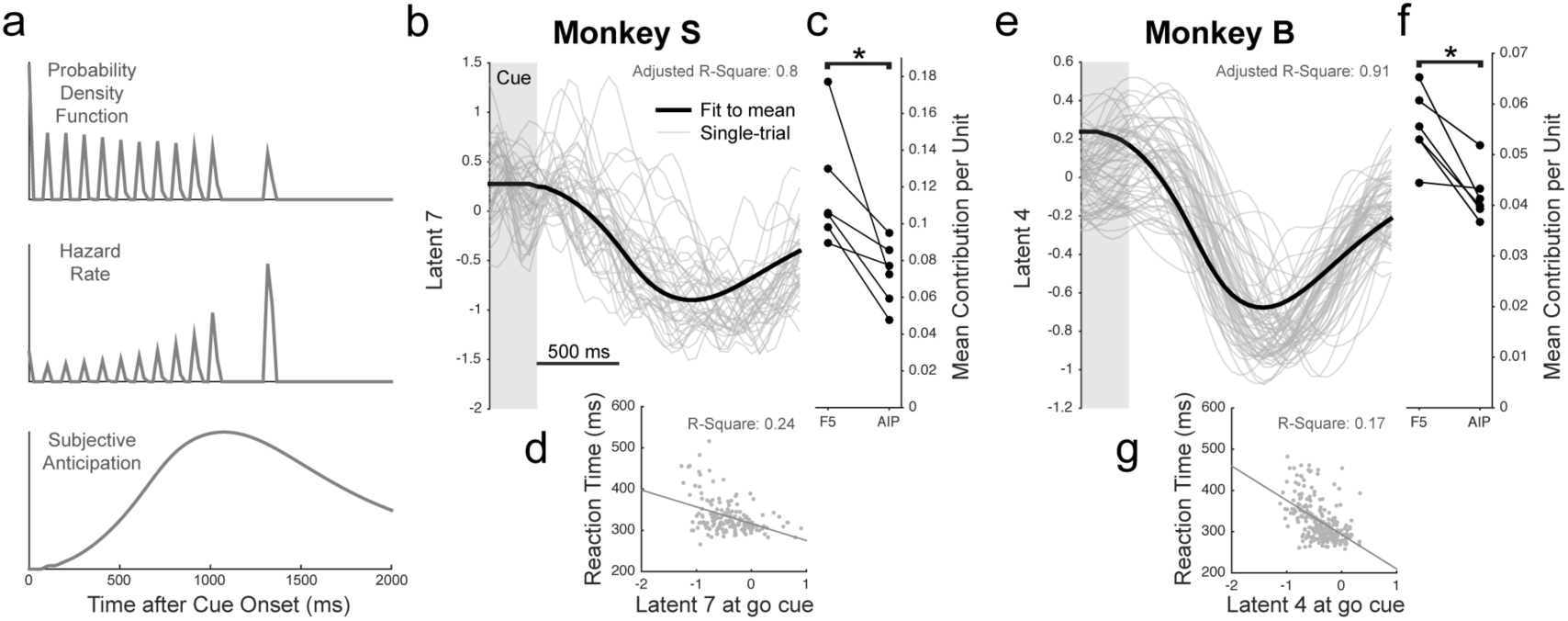
Representation of subjective anticipation across F5 and AIP. (a) Illustration of the probability of a go cue at all times during the delay, the hazard rate (Eq. 1), and the subjective anticipation function (Eq. 2 substituted into Eq. 1). (b) subjective anticipation (Eq. 3) fit to an example latent dimension during the no-movement condition (session S2). (c) Mean contribution per unit in each area to the best latent dimension of each data set. Stars indicate a significant difference (Wilcoxon sign-rank test, p < 0.001). (d) Example latent dimension at go cue correlated with single-trial reaction time for delays of at least 800 ms. (e-g) Same as (b-d) for monkey B (session B4).

When comparing the mean contribution per unit (weight in GPFA loading matrix) between areas across data sets to the best fitting latent dimensions, F5 clearly contributes more (Figure 7c,f, Wilcoxon signed-rank test, p < 0.001), with an average of 1.5 times the contribution per neuron, supporting the finding that F5 memory activity was much more dynamic. On average across data sets, the best fitting latent dimension explained the 4^th^ most variance of the 10 dimensions extracted for each data set, corresponding to on average 11% variance explained.

Interestingly, activity on single trials in the ideal latent dimensions at the go cue was correlated with reaction time (Figure 7d,g; trials with a delay of at least 800 ms), with a mean R-square of 0.17 in monkey S and 0.16 in monkey B, similar to results obtained in F5 with other state space methods (Michaels et al., 2015). For this analysis only the causal portion of all GPFA smoothing kernels were used so that activity at the go cue conservatively reflected only past spikes. Given that the activity in this latent dimension is predictive of reaction time, does being closer or farther away from the movement state predict reaction time in a consistent way? When the absolute difference between the go cue activity and mean activity during movement initiation (100 ms before movement onset) was correlated with reaction time, 11 out of 12 data sets produced a positive correlation (mean R-square of 0.1), providing evidence that being closer to the movement initiation state on a given trial led to shorter reaction times.

### Clustering of immediate and withheld movements from memory

In the population visualization in Figure 3 we saw that the trajectories of short and long delays formed two distinct clusters leading up to movement onset. To visualize the clustering for example data sets in F5, we plotted the activity of all linearly spaced delays (0-1000 ms) of a single grip type around movement onset in an example latent dimension (Fig 8a). Looking specifically at around 100 ms before movement onset, trajectories from the conditions with a delay of 0-400/500 ms and from the conditions with a delay of 400/500-1000 ms seem to form two clusters. This effect is also present in AIP, where trajectories deflect into two distinct groups in a similar fashion (Figure 9).

**Figure 8.**
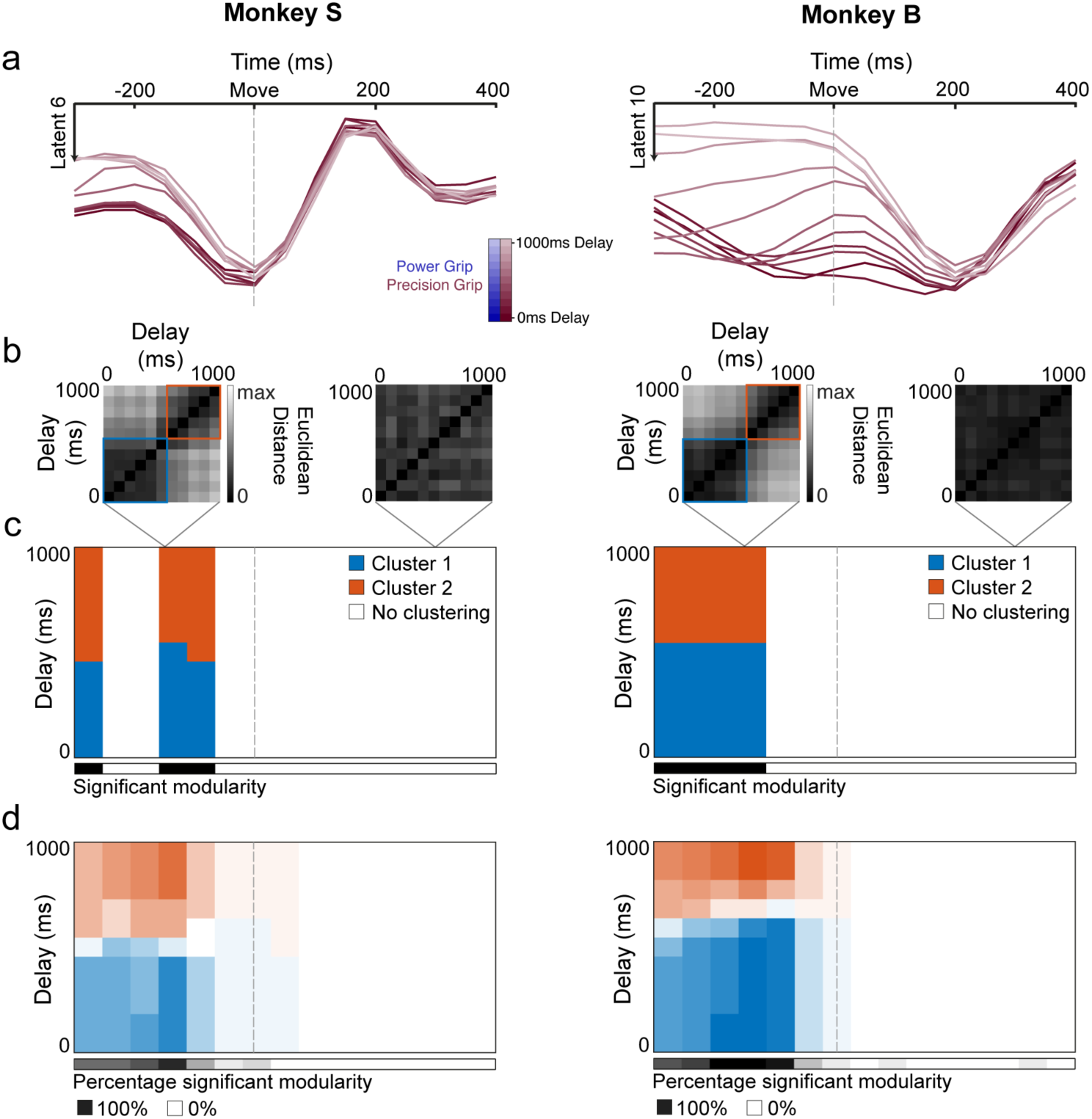
Clustering of movement initiation activity in F5. **(a)** Example latent projection population activity in F5 over all linearly spaced delays (0-1000 ms) for precision grip trials for an example data set from each monkey (sessions S4, B2), aligned to movement onset. (b) Euclidean distance between all pairs of delays in the full latent space for two example time points of the example data set including identified clustering using a clustering analysis that finds community structure (Materials and Methods). (c) Clusters identified in the distance matrices over time (in steps of 50 ms) for the example data set. Black significance bar shows time points where the modularity statistic exceeded chance level (permutation test, p < 0.01). (d) Same analysis as (c) averaged over all data sets and grip types (6 data sets x 2 grip types).

**Figure 9.**
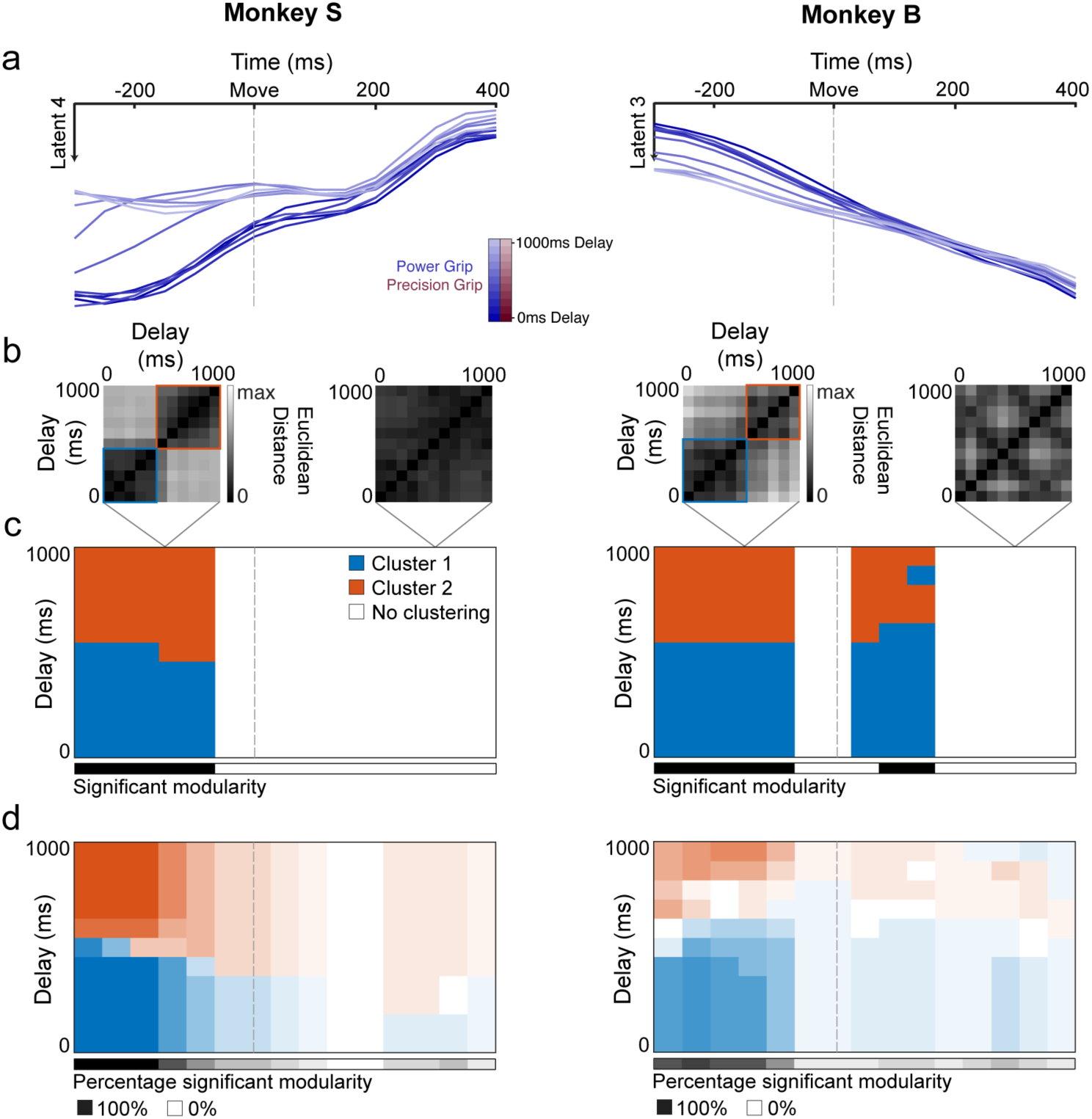
Clustering of movement initiation activity in AIP. Same Layout as Figure 8. (a) Example latent projection population activity in AIP over all linearly spaced delays (0-1000 ms) for precision grip trials for an example data set from each monkey (S3, B4), aligned to movement onset. (b) Euclidean distance between all pairs of delays in the full latent space for two example time points of the example data set including identified clustering using a clustering analysis that finds community structure (Materials and Methods). (c) Clusters identified in the distance matrices over time (in steps of 50 ms) for the example data set. Black significance bar shows time points where the modularity statistic exceeded chance level (permutation test, p < 0.01). (d) Same analysis as (c) averaged over all data sets and grip types (6 data sets x 2 grip types).

To quantify clustering at the population level, we calculated the Euclidean distance between all pairs of delay lengths for each grip type separately in the space of all latent dimensions (Figure 8b) and looked for clusters in the distance matrices without assuming clustering a priori (Materials and Methods). Two clusters were identified for the example data set (Figure 8c), showing a split around the 400-500 ms delay point that lasts until shortly before movement onset (permutation test, p < 0.01; Materials and Methods). This pattern was very similar over all data sets (Figure 8d, Figure 9d), did not differ between grip types, and was present in both areas and monkeys, indicating that the state change that occurs between short and long delays spans both the frontal and parietal lobes.

Clustering is not likely due to different movement kinematics, since the movement times were nearly identical for all delay lengths (Figure 1d), especially for monkey S. However, since the time of movement onset is determined by the monkey’s behavior, the time that has elapsed since the visual grip cue was presented could introduce a potential confound. Yet, differences in how long ago the grip cue was presented is unlikely to explain the two clusters, since repeating the same clustering analysis on the behavioral data, i.e. the mean time between cue presentation and movement onset for all delays, does not produce significant clustering for either grip type (permutation test, Precision grip: p = 0.97, Power grip: p = 0.97). These controls suggest that the separation of the neural trajectories into two distinct clusters reflects a robust effect of delay length in F5 and AIP.

## Discussion

To systematically probe the interplay between planning and movement in the grasping network, we recorded neural populations in premotor area F5 and parietal area AIP while two macaque monkeys performed a delayed grasping task with 12 distinct preparation times (0-1300 ms). Firstly, the initial part (∼300 ms) of the neural space traversed was the same for all delays, but was grip specific, providing evidence that this activity was an unavoidable part of preparing the correct movement. Next, population activity shifted into a separate state that was not achieved during short delays. The memory state was more dynamic in F5 than in AIP, tracking subjective movement anticipation over time. Lastly, activity during movement initiation formed two distinct clusters during movement initiation, demonstrating a network-wide shift when movements need to be withheld. Our findings reinforce the notion that more global aspects of movements, such as the movement plan, as well as dynamic aspects, such as cue anticipation, can be well extracted at the population level.

As shown in Figure 4, separation between the neural trajectories occurred more than 200 ms earlier between the two grips than between long and short delays. This novel result indicates that while grip information is swiftly encoded in F5 and AIP following the cue, responses to the go cue are delayed at least 200 ms relative to the grip information in order to facilitate the completion of the motor plan, after which areas of the state space traversed by longer delays are not strictly necessary to produce successful movements, similar to the results of Ames et al. (2014) in dorsal premotor cortex (PMd).

In F5 the memory period activity did not congregate in a specific region of the state space, a feature of F5 never before observed to our knowledge. This finding differs to the results of Ames et al. (2014) in nearby PMd, who postulated that delay period activity may act as an attractor state into which all trials would congregate given enough preparation time. It is possible that PMd activity would be more dynamic if an experimental design with a memory period were utilized, a point supported by studies showing that activity from some sub-regions of premotor cortex can encode prior knowledge of when events are likely to occur (Mauritz and Wise, 1986; Carnevale et al., 2015). However, given current evidence our results support the notion that strongly dynamic memory period activity is a unique feature of F5.

It could be that the temporal dynamics during the memory period are a result of an internalized representation of the likelihood of task events occurring at specific times throughout the memory period, known as hazard rate and previously observed in the lateral intraparietal cortex (LIP) (Leon and Shadlen, 2003; Janssen and Shadlen, 2005). We observed significant fits of latent dimensions to the subjective anticipation rate across both areas, although F5 contributed significantly more to this activity. Furthermore, activity in these dimensions was predictive of reaction time, supporting the role of this activity in increasing or decreasing sensitivity to an external stimulus.

Time dependence has been identified in prefrontal areas (Genovesio et al., 2006), and increasing literature suggesting that time keeping is an intrinsic property of all neural networks (for a review see Goel and Buonomano, 2014), as well as a feature of some sub-cortical areas (Gouvêa et al., 2015). A mechanistic explanation for the dynamics observed during the memory period could be that recurrent networks of neurons in these areas generate temporal dynamics similar to a time code. The strongest evidence for this view comes from a recent study in which the presence or absence of a sensory stimulus on a given trial had to be reported (Carnevale et al., 2015). The authors found that the neural state space of premotor cortex evolved over the course of the trial and was more sensitive to incoming sensory information during the fixed window that the monkeys knew would or would not contain the stimulus. Importantly, Carnevale et al. (2015) showed that a recurrent neural network model trained for optimal response sensitivity well explained the behavior of the monkey. A number of recent studies have shown that timing is a robust feature of chaotic recurrent networks (Buonomano and Laje, 2010; Laje et al., 2013; Goudar and Buonomano, 2014), suggesting that F5 may be able to track the course of time internally and use this information to predict when an action is likely to be required. Furthermore, even though activity continues to change throughout memory, a stable representation of the desired action remains at the population level (Druckmann and Chklovskii, 2012), consistent with the constant separation between grip types observed in some latent dimensions (Figure 3).

One of the most striking features in both areas, but especially in F5, was that the population activity of a single grip type was highly variable at the time of go cue, yet converged rapidly leading up to movement onset (Figure 3, Figure 8). We propose that the broadly tuned nature of activity at the go cue provides the motor system with a large flexibility in movement initiation. Similar to the dynamics observed during the memory period, it could be that once movement is triggered, recurrent networks of neurons within these areas rapidly reduce variability within particular regions of the neural space in order to ensure correct muscle activation during initiation (Sussillo et al., 2015; Michaels et al., 2016). Under this framework, selecting between multiple movement plans would only require the neural population to be within a general region of activity. Such a framework is also in line with the finding that preparatory activity in PMd/M1 projects into the null-space of upper limb muscles and transitions into the potent-space during movement (Kaufman et al., 2014), as this transition likely takes place during movement initiation when variability between movement plans is heavily reduced (Elsayed et al., 2016). Once movement is initiated, activity would fall onto a common trajectory unique to each action plan. Future work must tackle the question of to what degree local circuit features or extrinsic inputs can account for the rapid decrease in trial-to-trial variability taking place before movement execution.

While variability decreased leading up to movement onset, trajectories clustered into two distinct groups splitting between delay conditions less than or greater than 400-500 ms (Figure 8, Figure 9). Given that full preparation likely takes ∼400 ms, evidenced by the leveling of the RT curve after ∼400 ms (Figure 1d), the two clusters could correspond to movements executed ‘as fast as possible’ and movements executed from memory where the monkey must first wait for the go signal. Our results indicate that shifting between immediate movements and withheld movements from memory may cause a state shift in the fronto-parietal network that produces the two clusters during movement initiation. Once the state has been changed, the trajectories continue to cluster for the entirety of movement initiation (up to movement onset). Specifically, the underlying cause of the shift is likely the transition from reactive to proactive control, i.e., the increased ability to properly anticipate a go cue after sufficient preparation times (Braver, 2012). This sensitivity to task timing is inherent in highly trained tasks, and has been shown in supplementary motor area (SMA; Chen et al., 2010) and medial frontal cortex (Stuphorn and Emeric, 2012). Execution of timed behavior is reduced in humans with SMA lesions (Halsband et al., 1993) and supports our findings, since F5 is especially connected to the pre-SMA (Luppino et al., 1993).

It remains a possibility that systematic differences in hand-shaping latencies or final posture between different delay lengths could contribute to the observed clustering. However, clustering of delay conditions was almost non-existent after movement onset, especially in F5, making differences in final posture improbable. Although differences in hand-shaping during movement cannot be ruled out, the extreme similarity in movement times between delays (Results), especially for monkey S, make this possibility unlikely.

Given that the current task also involved a large reaching component, reach planning is likely a significant part of the observed activity. Still, the presence of grip type tuning in all epochs (Results), as well as previous research employing a grasp-only task (Hepp-Reymond et al., 1994) and a grasp-reach dissociation task (Lehmann and Scherberger, 2013), indicates that F5 encodes grasping quite independently of reaching. Furthermore, reversibly inactivating F5 (Fogassi et al., 2001) or AIP (Gallese et al., 1994) selectively impairs hand-shaping and not reaching, providing evidence that our results are an accurate representation of the grasping network.

In summary, our results provide novel insights building on delayed reaching and grasping literature in premotor (Cisek et al., 2003; Lucchetti et al., 2005; Fluet et al., 2010) and parietal cortex (Murata et al., 1996; Snyder et al., 2006; Baumann et al., 2009). We show that dissociation of global and dynamic aspects of movement, such as the movement plan and the anticipation over time, respectively, can be coherently extracted at the level of neural populations and allow for comparison and dissociation between interacting cortical areas.

## Acknowledgements

We would like to thank Natalie Bobb, Ricarda Lbik, and Matthias Dörge for technical assistance, and Roman Eppinger for preliminary analysis.

## Notes

Conflict of Interest: The authors declare no competing financial interests.

## References

Afshar A, Santhanam G, Yu BM, Ryu SI, Sahani M, Shenoy KV (2011) Single-trial neural correlates of arm movement preparation. Neuron 71:555–564.

Ames KC, Ryu SI, Shenoy KV (2014) Neural Dynamics of Reaching following Incorrect or Absent Motor Preparation. Neuron 81:438–451.

Baumann MA, Fluet M-C, Scherberger H (2009) Context-specific grasp movement representation in the macaque anterior intraparietal area. J Neurosci 29:6436–6448.

Braver TS (2012) The variable nature of cognitive control: a dual mechanisms framework. Trends in Cognitive Sciences 16:106–113.

Buonomano DV, Laje R (2010) Population clocks: motor timing with neural dynamics. Trends in Cognitive Sciences 14:520–527.

Carnevale F, de Lafuente V, Romo R, Barak O, Parga N (2015) Dynamic Control of Response Criterion in Premotor Cortex during Perceptual Detection under Temporal Uncertainty. Neuron 86:1067–1077.

Chen X, Scangos KW, Stuphorn V (2010) Supplementary motor area exerts proactive and reactive control of arm movements. J Neurosci 30:14657– 14675.

Churchland MM, Cunningham JP, Kaufman MT, Foster JD, Nuyujukian P, Ryu SI, Shenoy KV (2012) Neural population dynamics during reaching. Nature 487:51–56.

Churchland MM, Shenoy KV (2007) Delay of movement caused by disruption of cortical preparatory activity. J Neurophysiol 97:348–359.

Churchland MM, Yu BM, Ryu SI, Santhanam G, Shenoy KV (2006) Neural variability in premotor cortex provides a signature of motor preparation. J Neurosci 26:3697–3712.

Cisek P, Crammond DJ, Kalaska JF (2003) Neural activity in primary motor and dorsal premotor cortex in reaching tasks with the contralateral versus ipsilateral arm. J Neurophysiol 89:922–942.

Crammond DJ, Kalaska JF (2000) Prior information in motor and premotor cortex: activity during the delay period and effect on pre-movement activity. J Neurophysiol 84:986–1005.

Cunningham JP, Yu BM (2014) Dimensionality reduction for large-scale neural recordings. Nat Neurosci 17:1500–1509.

Dann B, Michaels JA, Schaffelhofer S, Scherberger H (2016) Uniting functional network topology and oscillations in the fronto-parietal single unit network of behaving primates. eLife.

Day BL, Rothwell JC, Thompson PD, Maertens de Noordhout A, Nakashima K, Shannon K, Marsden CD (1989) Delay in the execution of voluntary movement by electrical or magnetic brain stimulation in intact man. Evidence for the storage of motor programs in the brain. Brain 112 (Pt 3):649–663.

Druckmann S, Chklovskii DB (2012) Neuronal circuits underlying persistent representations despite time varying activity. Curr Biol 22:2095–2103.

Elsayed GF, Lara AH, Kaufman MT, Churchland MM, Cunningham JP (2016) Reorganization between preparatory and movement population responses in motor cortex. Nat Commun 7:13239.

Fluet M-C, Baumann MA, Scherberger H (2010) Context-specific grasp movement representation in macaque ventral premotor cortex. J Neurosci 30:15175–15184.

Fogassi L, Gallese V, Buccino G, Craighero L, Fadiga L, Rizzolatti G (2001) Cortical mechanism for the visual guidance of hand grasping movements in the monkey: A reversible inactivation study. Brain 124:571–586.

Gallese V, Murata A, Kaseda M, Niki N, Sakata H (1994) Deficit of hand preshaping after muscimol injection in monkey parietal cortex. Neuroreport 5:1525–1529.

Genovesio A, Tsujimoto S, Wise SP (2006) Neuronal activity related to elapsed time in prefrontal cortex. J Neurophysiol 95:3281–3285.

Gerits A, Farivar R, Rosen BR, Wald LL, Boyden ES, Vanduffel W (2012) Optogenetically induced behavioral and functional network changes in primates. Curr Biol 22:1722–1726.

Ghez C, Favilla M, Ghilardi MF, Gordon J, Bermejo R, Pullman S (1997) Discrete and continuous planning of hand movements and isometric force trajectories. Exp Brain Res 115:217–233.

Goel A, Buonomano DV (2014) Timing as an intrinsic property of neural networks: evidence from in vivo and in vitro experiments. Philos Trans R Soc Lond, B, Biol Sci 369:20120460.

Goudar V, Buonomano DV (2014) Useful dynamic regimes emerge in recurrent networks. Nat Neurosci 17:487–489.

Gouvêa TS, Monteiro T, Motiwala A, Soares S (2015) Striatal dynamics explain duration judgments. eLife.

Gozani SN, Miller JP (1994) Optimal discrimination and classification of neuronal action potential waveforms from multiunit, multichannel recordings using software-based linear filters. IEEE Trans Biomed Eng 41:358–372.

Halsband U, Ito N, Tanji J, Freund HJ (1993) The role of premotor cortex and the supplementary motor area in the temporal control of movement in man. Brain 116 (Pt 1):243–266.

Hepp-Reymond MC, Hüsler EJ, Maier MA, Qi HX (1994) Force-related neuronal activity in two regions of the primate ventral premotor cortex. Can J Physiol Pharmacol 72:571–579.

Janssen P, Shadlen MN (2005) A representation of the hazard rate of elapsed time in macaque area LIP. Nat Neurosci 8:234–241.

Kaufman MT, Churchland MM, Ryu SI, Shenoy KV (2014) Cortical activity in the null space: permitting preparation without movement. Nat Neurosci 17:440– 448.

Kaufman MT, Seely JS, Sussillo D, Ryu SI, Shenoy KV, Churchland MM (2016) The Largest Response Component in the Motor Cortex Reflects Movement Timing but Not Movement Type. eNeuro 3.

Kraskov A, Dancause N, Quallo MM, Shepherd S, Lemon RN (2009) Corticospinal Neurons in Macaque Ventral Premotor Cortex with Mirror Properties: A Potential Mechanism for Action Suppression? Neuron 64:922–930.

Kutas M, Donchin E (1974) Studies of squeezing: handedness, responding hand, response force, and asymmetry of readiness potential. Science 186:545–548.

Laje R, Buonomano DV, Buonomano DV (2013) Robust timing and motor patterns by taming chaos in recurrent neural networks. Nat Neurosci 16:925–933.

Lehmann SJ, Scherberger H (2013) Reach and gaze representations in macaque parietal and premotor grasp areas. J Neurosci 33:7038–7049.

Leon MI, Shadlen MN (2003) Representation of time by neurons in the posterior parietal cortex of the macaque. Neuron 38:317–327.

Lucchetti C, Ulrici A, Bon L (2005) Dorsal premotor areas of nonhuman primate: functional flexibility in time domain. Eur J Appl Physiol 95:121–130.

Luppino GG, Luppino G, Matelli MM, Matelli M, Camarda R, Camarda RR, Rizzolatti G, Rizzolatti GG (1993) Corticocortical connections of area F3 (SMA-proper) and area F6 (pre-SMA) in the macaque monkey. J Comp Neurol 338:114–140.

Maris E, Oostenveld R (2007) Nonparametric statistical testing of EEG- and MEG-data. J Neurosci Methods 164:177–190.

Mauritz KH, Wise SP (1986) Premotor cortex of the rhesus monkey: neuronal activity in anticipation of predictable environmental events. Exp Brain Res 61:229–244.

Menz VK, Schaffelhofer S, Scherberger H (2015) Representation of continuous hand and arm movements in macaque areas M1, F5, and AIP: a comparative decoding study. J Neural Eng 12:056016.

Michaels JA, Dann B, Intveld RW, Scherberger H (2015) Predicting Reaction Time from the Neural State Space of the Premotor and Parietal Grasping Network. J Neurosci 35:11415–11432.

Michaels JA, Dann B, Scherberger H (2016) Neural Population Dynamics during Reaching Are Better Explained by a Dynamical System than Representational Tuning. PLoS Comput Biol 12:e1005175.

Murata A, Gallese V, Kaseda M, Sakata H (1996) Parietal neurons related to memory-guided hand manipulation. J Neurophysiol 75:2180–2186.

Musial PG, Baker SN, Gerstein GL, King EA, Keating JG (2002) Signal-to-noise ratio improvement in multiple electrode recording. J Neurosci Methods 115:29–43.

National Research Council (2003) Guidelines for the Care and Use of Mammals in Neuroscience and Behavioral Research. Washington (DC): National Academies Press (US).

Newman MEJ (2004) Fast algorithm for detecting community structure in networks. Phys Rev E 69:066133.

Quiroga RQ, Nadasdy Z, Ben-Shaul Y (2004) Unsupervised spike detection and sorting with wavelets and superparamagnetic clustering. Neural Comput 16:1661–1687.

Reichardt J, Bornholdt S (2006) Statistical mechanics of community detection. Phys Rev E 74:016110.

Riehle A, Requin J (1989) Monkey primary motor and premotor cortex: single-cell activity related to prior information about direction and extent of an intended movement. J Neurophysiol 61:534–549.

Rosenbaum DA (1980) Human movement initiation: specification of arm, direction, and extent. J Exp Psychol Gen 109:444–474.

Schaffelhofer S, Agudelo-Toro A, Scherberger H (2015) Decoding a wide range of hand configurations from macaque motor, premotor, and parietal cortices. J Neurosci 35:1068–1081.

Schaffelhofer S, Scherberger H (2016) Object vision to hand action in macaque parietal, premotor, and motor cortices. eLife 5:6436.

Snyder LH, Dickinson AR, Calton JL (2006) Preparatory delay activity in the monkey parietal reach region predicts reach reaction times. J Neurosci 26:10091–10099.

Stuphorn V, Emeric EE (2012) Proactive and reactive control by the medial frontal cortex. Front Neuroeng 5:9.

Sussillo D, Churchland MM, Kaufman MT, Shenoy KV (2015) A neural network that finds a naturalistic solution for the production of muscle activity. Nat Neurosci 18:1025–1033.

Townsend BR, Subasi E, Scherberger H (2011) Grasp movement decoding from premotor and parietal cortex. J Neurosci 31:14386–14398.

Wise SP (1985) The primate premotor cortex: past, present, and preparatory. Annu Rev Neurosci 8:1–19.

Wise SP, Kurata K (1989) Set-Related Activity in the Premotor Cortex of Rhesus Monkeys: Effect of Triggering Cues and Relatively Long Delay Intervals. Somatosens Mot Res 6:455–476.

Yu BM, Cunningham JP, Santhanam G, Ryu SI, Shenoy KV, Sahani M (2009) Gaussian-process factor analysis for low-dimensional single-trial analysis of neural population activity. J Neurophysiol 102:614–635.

